## Supplementary information for "Sustainable reference points for multispecies coral reef fisheries"

14th of September 2022

**This PDF file includes:**

Figs. S1 to S8
Tables S1 to S3
Supplementary Information 1:
- Supplementary information 1.1: Sensitivity analyses to the choice of surplus production model (Fig. SI1-1; Table SI1-1)
- Supplementary information 1.2: Comparison with previous reference point estimates (Table SI1-2)
- Supplementary information 1.3: Trying to parameterize exports
- Supplementary information 1.4: Species intrinsic growth rates vs. community growth rates (Fig. SI1-2)
- Supplementary information 1.5: Model fits (Fig. SI1-3-4)
- Supplementary information 1.6: Checking data inclusion (Fig. SI1-5-9)
- Supplementary information 1.7: Study workflow (Fig SI1-10)

Supplementary Information 2: Principled Bayesian workflow (Fig. SI2-1-12; Table SI2-1-2)

**Other Supplementary Materials for this manuscript include the following:**

Data and code: Data and code used for this paper will be available from GitHub (https://github.com/ JZamborain-Mason/ZamborainMasonetal2022_Sustainability).

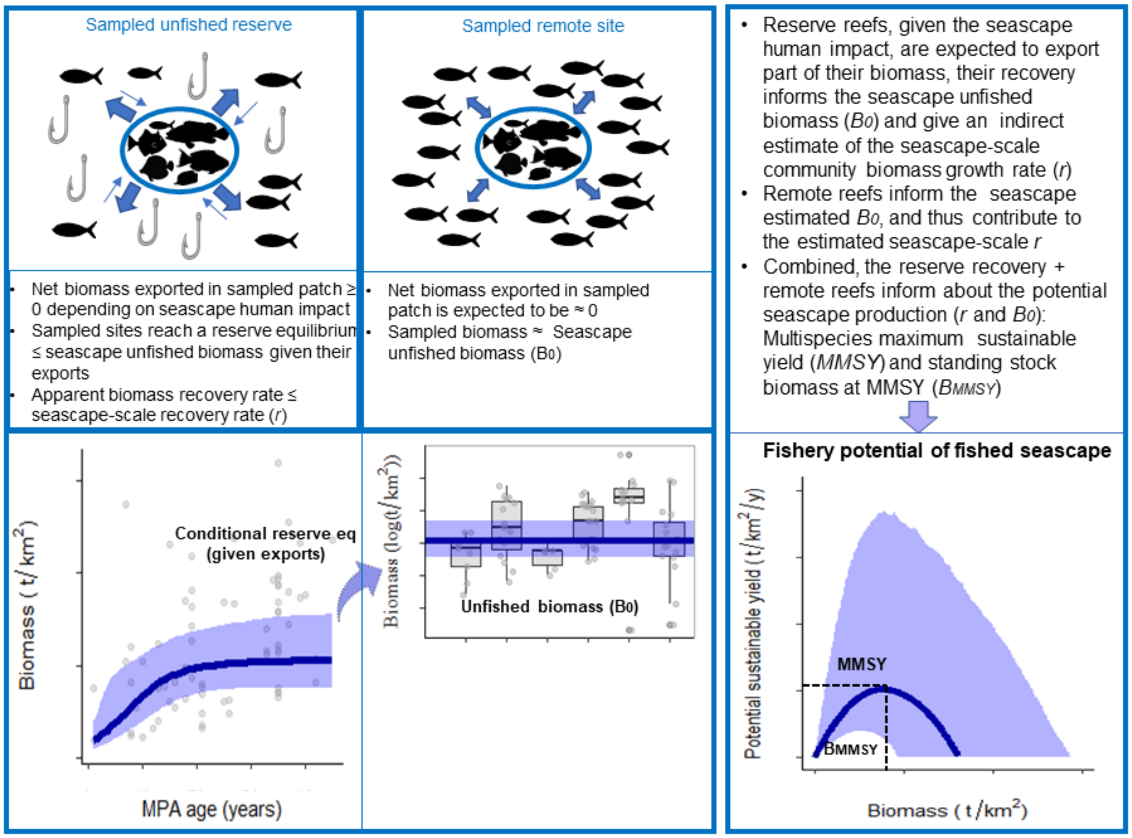

**Fig. S1| Diagram explaining how reserve and remote reefs inform (seascape-scale) sustainable reference points.**

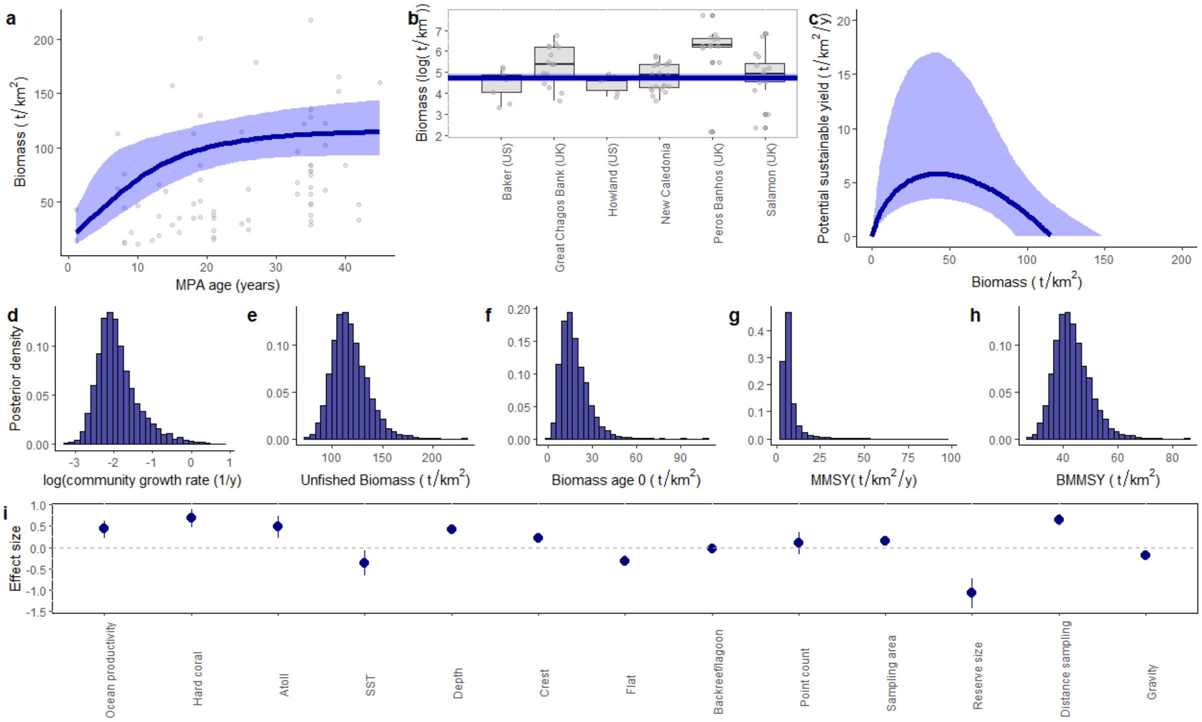

**Fig. S2| Model fit and parameter posterior distributions from the best-fit joint Bayesian MMSY and BMMSY benchmark model for average and most common environmental conditions (e.g., non-atolls).** (a) Fitted reserve trajectory. Each dot represents the median estimated biomass at a reserve site adjusted for methodological covariates. (b) Estimated unfished biomass over the estimated biomass at remote locations adjusted for methodological covariates. y axes is log-transformed to aid clarity of distributions. (c) Estimated surplus production curve. (d-g) Posterior parameter density distributions for average and most common conditions and zero human impact: (d) community growth rate (0.14 [0.08 - 0.31] 1/y; (posterior median [90% posterior uncertainty intervals]), (e) unfished biomass (115.6 [97.5 - 140.6] $t/km^2$), (f) biomass at reserve age 0 (16.3 [8.2 – 30.0] $t/km^2$), (g) estimated MMSY for average environmental conditions (5.8[3.8 - 12.3] $t/km^2/y$), and (h) BMMSY for average environmental conditions (42.5 [35.9-51.7] $t/km^2$). (i) Posterior effect sizes -median and 90% uncertainty intervals- of the covariates included in the model. Environmental covariates were a function of site's estimated unfished biomass, sampling covariates were a function of observed biomass, and reserve size was only used in the reserve subcomponent part of the model. In (a), (b) and (c) the line represents the Bayesian posterior median values and light polygons represent the 90 % uncertainty intervals.

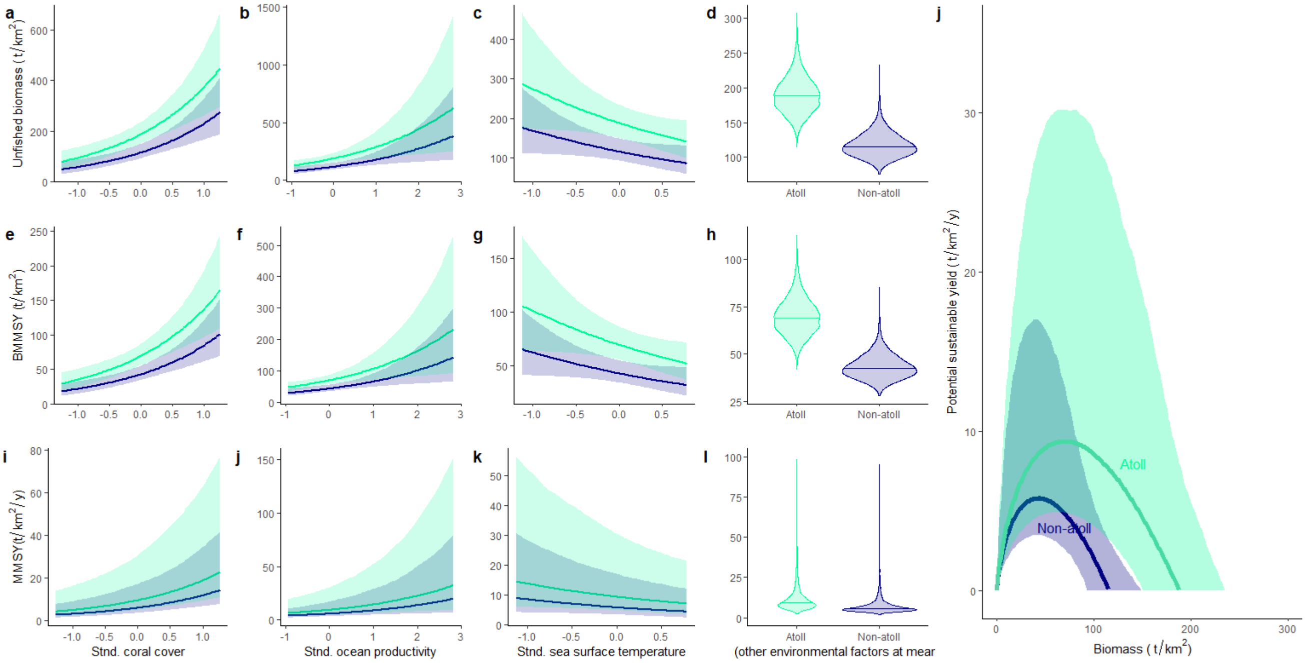

**Fig. S3| Estimated effect of environmental covariates on sustainable reference points.** Simulations are from model posteriors maintaining all other covariates at most common categories and average conditions. Except for d, h, l, the line represents the median and the polygons the 90% uncertainty intervals. In d, h, l the entire distribution is plotted. Note that for standardized coral cover, ocean productivity and SST, zero represents the average of our sampled reefs.

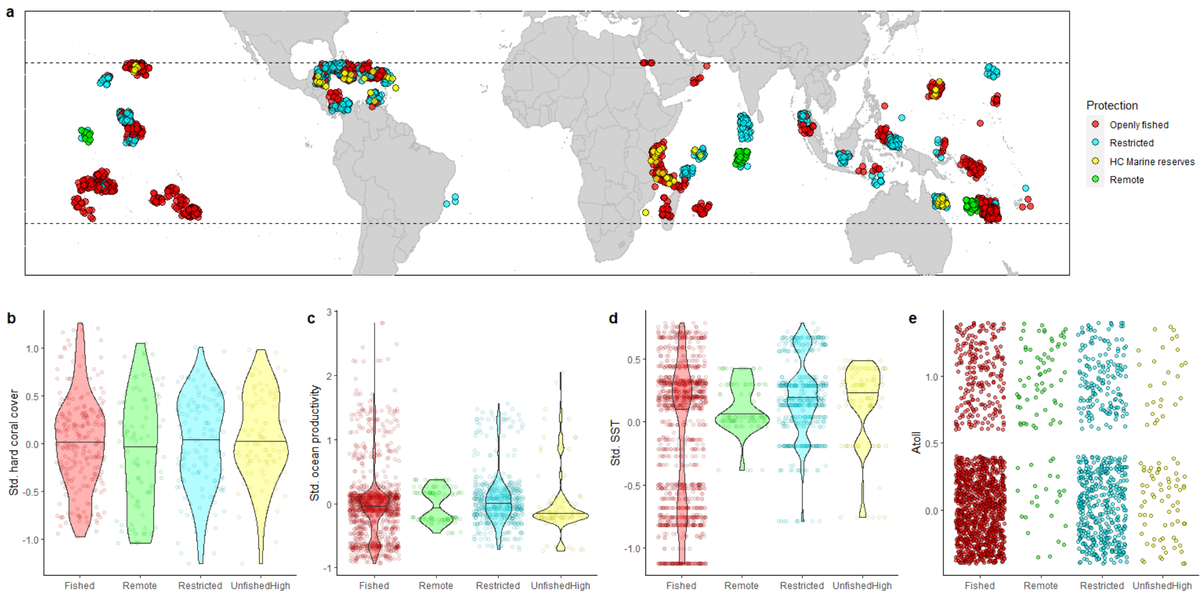

**Fig. S4| Sites used in this study.** (a) Map of our sampled sites used to estimate sustainable reference points (High Compliance marine reserve and Remote reefs) and assess the status of fished coral reef fish stocks (Restricted, and Openly Fished). Points are slightly jittered for clarity. (b-e) Environmental information for each site (standardized) separated by protection category. Note violin plots overlap indicating that there are no major differences in environmental factors among categories (e.g., reserve placement conditions are likely not biased, environmentally, in comparison to the rest of our data).

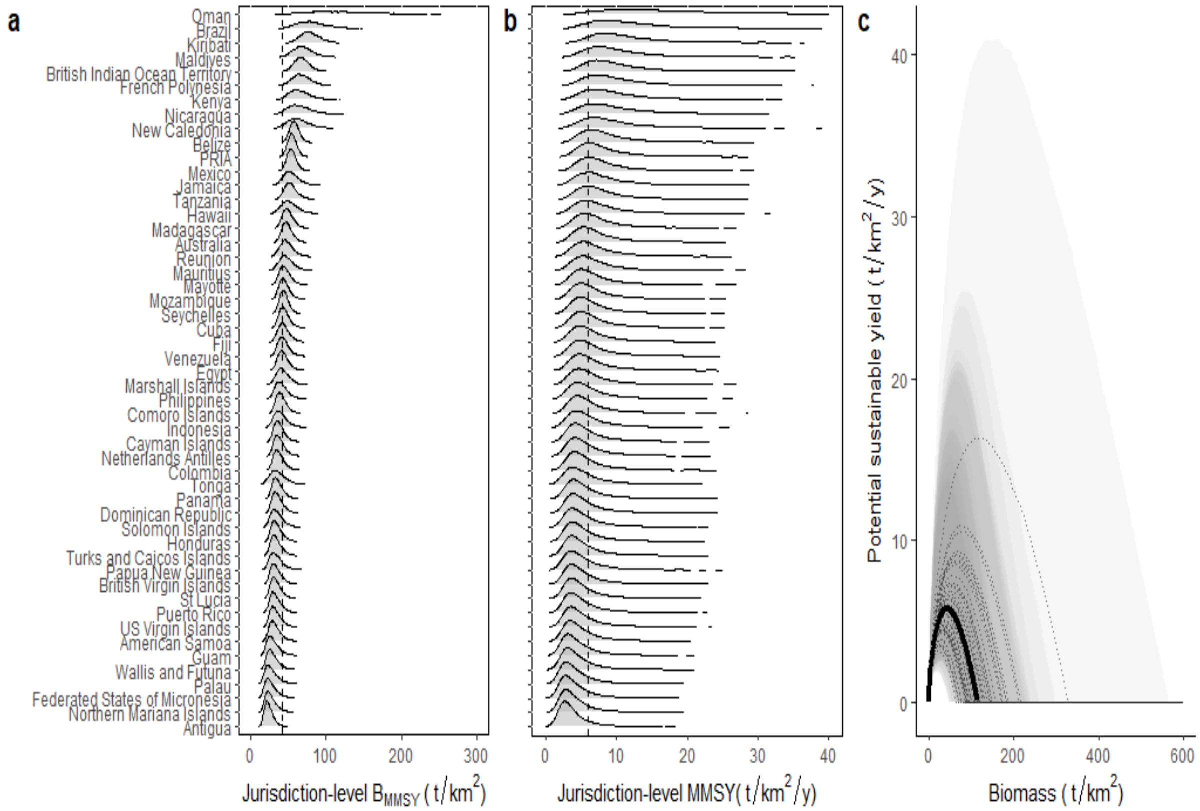

**Fig. S5| Jurisdiction-specific estimated sustainable reference points and surplus production curves (based on our sampled sites).** In a and b, distributions represent the posterior jurisdiction reference points (4000 samples). Vertical dashed lines represent the median for average and most common environmental conditions. In c grey lines represent the median jurisdiction-specific surplus production curve and the black line represents the median for average and most common environmental conditions. Light grey polygons are jurisdiction-specific 90% uncertainty intervals, with darker regions indicating higher overlap in uncertainty intervals. PRIA refers to Pacific Remote Islands and Atolls.

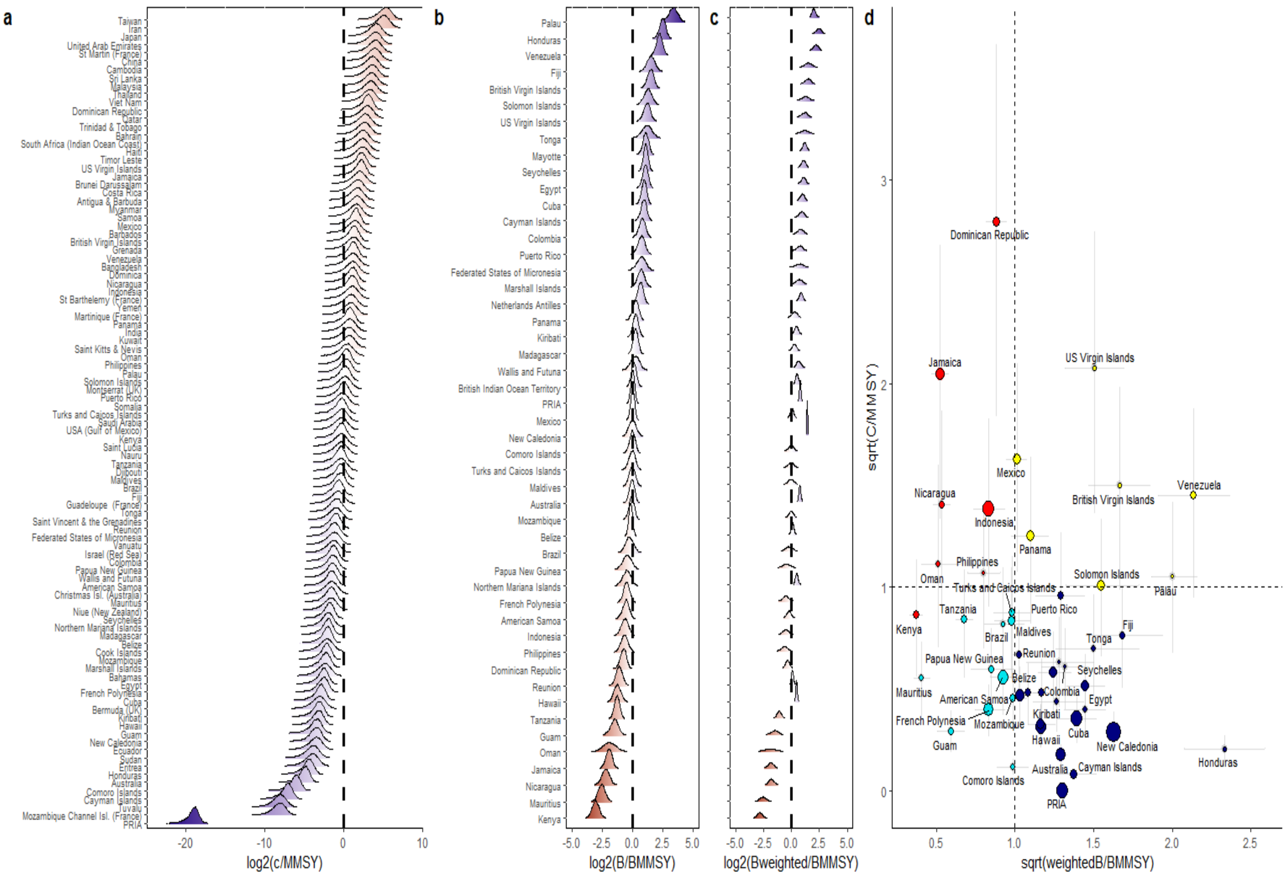

**Fig. S6| Status of fished (openly fished and/or restricted) reefs per jurisdiction.** (a-b) Fishing (C/MMSY) and biomass status ($B/B_{MMSY}$ or weighted by proportion of waters protected in jurisdiction: $B_{weighted}/B_{MMSY}$) for each jurisdiction given their MMSY and $B_{MMSY}$ distribution (i.e., showing the probabilities of catching above MMSY or having biomass values below $B_{MMSY}$ given the sustainable reference point distributions). Note that in (a) for jurisdictions that had catch estimates but without biomass data we used the jurisdiction combined posterior MMSY distribution. Vertical lines indicate where biomass and catch equal to $B_{MMSY}$ and MMSY, respectively. (c) Fishing vs. biomass (weighted) status plot for jurisdictions with both spatial-reconstructed catch and biomass data available (axes sqrt transformed). Color is based on fishery median status classification according to a jurisdiction's specific surplus curve: red= unsustainable, turquoise=recovering, yellow=warning, and navy blue= in good condition. Error bars represent the 0.9 quantiles of the jurisdiction posterior distributions. Size of the points is scaled according to the number of sites sampled for biomass**.** Note that estimated status is based on our sampled reefs open to extraction at the time of sampling. PRIA refers to Pacific Remote Islands and Atolls.

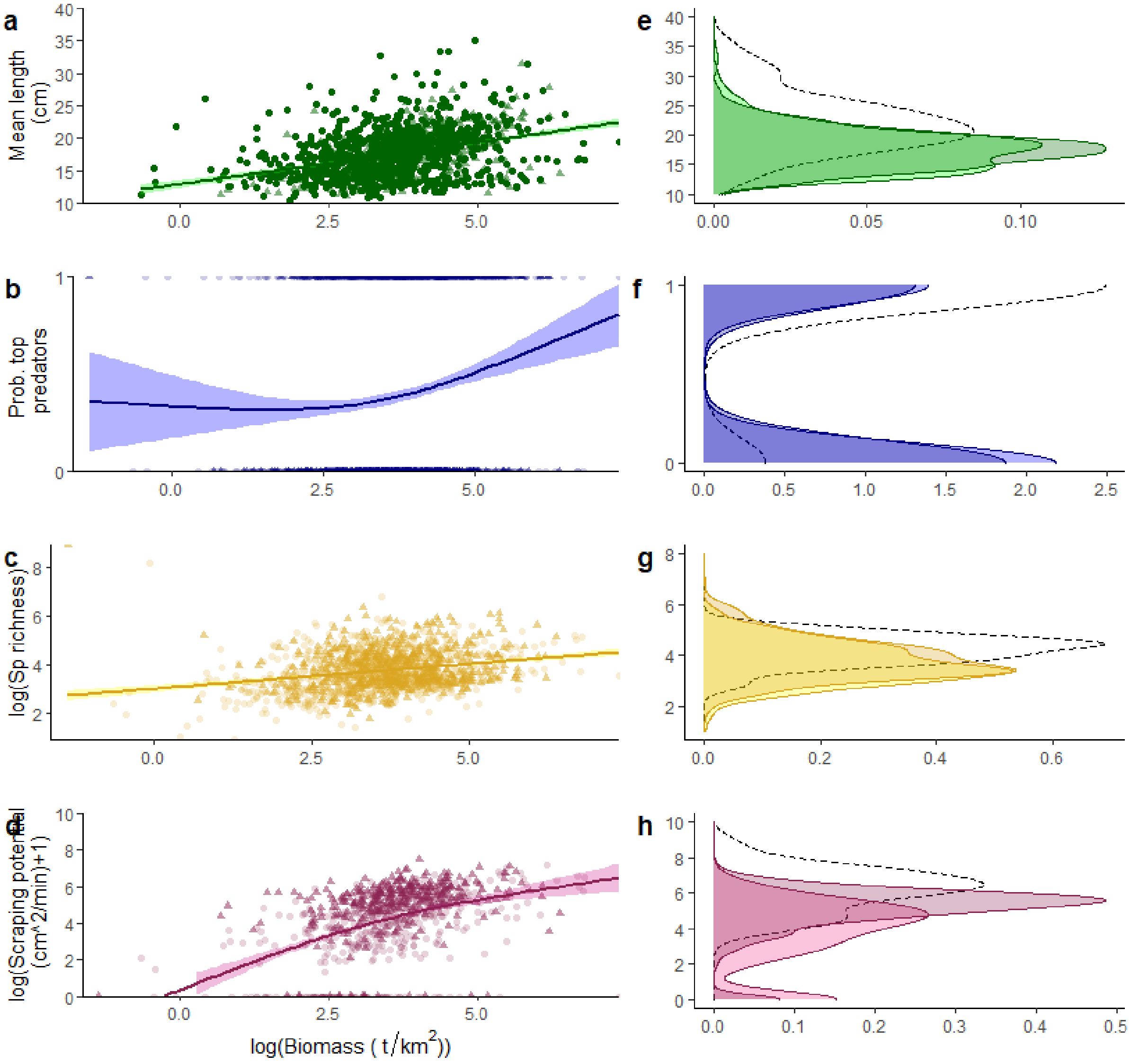

**Fig. S7| Relationship between site-specific biomass status and key ecosystem metrics and their distributions.** (a,c.e,g) Estimated relationship between marginalized biomass (on a logarithmic scale) and the four ecosystem metrics we examined; all corrected for environmental and sampling effects to represent average conditions and most common categories, to compare with the surplus curve. Solid line represents the best fit generalized additive model and polygons show 95% confidence intervals. Each point represents a single reef. Dark triangles represent reefs with some level of fishing restrictions and light points represent openly fished reefs. (b,d,f,g) Ecosystem metric distributions. Marginalized mean fish length, total richness (in a logarithmic scale), presence of top predators (density of 0's and 1's), and parrotfish scraping potential (in log +1 scale) of fish reefs separated by management (i.e., whether restrictions were in place (dark) or reefs were openly fished (light)). Dotted line is the overlaid distribution of observed response variables from our remote reefs (uninhabited >20 h away from human settlements).

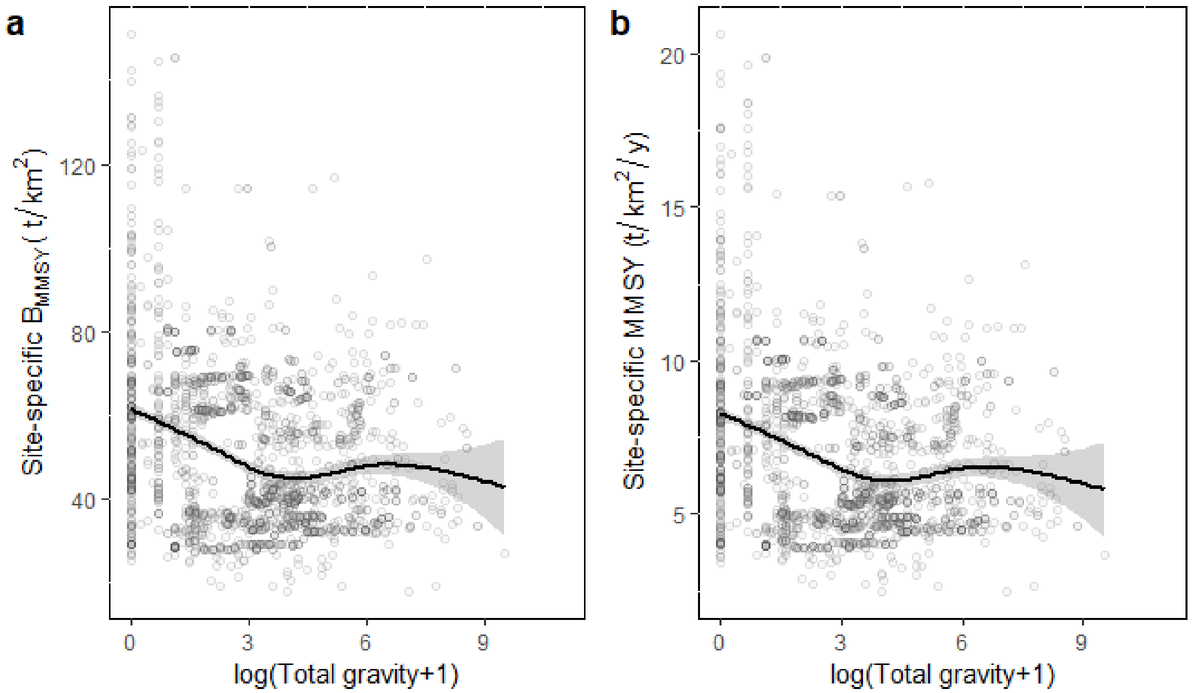

**Fig. S8| Relationship between site-specific $B_{MMSY}$ (a) and MMSY (b) and our metric of human impact (total gravity).** Posterior median estimated reference points for each site given their environmental conditions (based on our main model) as a function of total gravity- a proxy for human pressure on reefs. Each point represents a reef site, and trend line and polygon represent the mean and 95% confidence intervals of a generalized additive model fit to the relationship with total gravity (log+1 transformed).

**Table S1| Reef families included in our analyses.**

| Fish family | Common family name |
|---|---|
| **Acanthuridae** | Surgeonfishes |
| **Balistidae** | Triggerfishes |
| **Caesionidae** | Fusiliers |
| **Carangidae** | Jacks |
| **Chaetodontidae** | Butterflyfishes |
| **Cirrhitidae** | Hawkfishes |
| **Diodontidae** | Porcupinefishes |
| **Ephippidae** | Batfishes |
| **Haemulidae** | Sweetlips |
| **Kyphosidae** | Drummers |
| **Lethrinidae** | Emperors |
| **Lutjanidae** | Snappers |
| **Monacanthidae** | Filefishes |
| **Mullidae** | Goatfishes |
| **Nemipteridae** | Coral Breams |
| **Pinguipedidae** | Sandperches |
| **Pomacanthidae** | Angelfishes |
| **Labridae** | Wrasses and Parrotfishes |
| **Serranidae** | Groupers |
| **Siganidae** | Rabbitfishes |
| **Sparidae** | Porgies |
| **Sphyraenidae** | Barracudas |
| **Synodontidae** | Lizardfishes |
| **Tetraodontidae** | Pufferfishes |
| **Zanclidae** | Moorish Idol |

**Table S2| Model selection results for models in each section.** Values represent the expected log predictive density differences from model selection through leave-out-one cross validation.

| A) Reference point and assessment model | | | |
|---|---|---|---|
| **Model** | Elpd_diff | | Se_diff |
| **Full** | 0 | | 0 |
| **Null** | -493.1 | | 33.0 |

| B) Ecosystem metric models | | | | |
|---|---|---|---|---|
| **Model** | Parrotfish scraping potential | Top predator presence/absence | Mean length | Total fish species richness |
| **Full** | 0 | 0 | 0 | 0 |
| **Null** | -200.8 | -266.3 | -69.7 | -466.8 |

| C) Exploring alternate surplus production models | | |
|---|---|---|
| | Elpd_diff | Se_diff |
| **Gompertz-Fox (Full)** | 0 | 0 |
| **Graham-Schaefer** | -0.9 | 0.7 |
| **Pella-Tomlinson, 3** | -1.5 | 1.2 |
| **Pella-Tomlinson, 4** | -2.5 | 1.5 |

| D) Exploring different export parametrizations | | |
|---|---|---|
| | Elpd_diff | Se_diff |
| **No exports but modelling parameters with unstandardized gravity (Full)** | 0 | 0 |
| **Exports as a proportion of the community growth rate** | -67.7 | 12.5 |
| **Exports as a rate** | -68.4 | 12.5 |

| E) Fixing export proportions (as a function of community growth rate) | | |
|---|---|---|
| | Elpd_diff | Se_diff |
| **No exports but modelling parameters with unstandardized gravity (Full)** | 0 | 0 |
| **10%** | -66.4 | 12.5 |
| **15%** | -66.4 | 12.5 |
| **0%** | -66.6 | 12.5 |
| **30%** | -66.7 | 12.5 |
| **5%** | -66.7 | 12.5 |
| **20%** | -66.8 | 12.5 |
| **25%** | -67.1 | 12.5 |

**Table S3| Jurisdictions included in our status analyses.** "*" means information was available.

| Jurisdiction | Catch_data | Biomass_data | Number of available sampled sites |
|---|---|---|---|
| **American Samoa** | * | * | 104 |
| **Anguilla (UK)** | * | | |
| **Antigua & Barbuda** | * | * | 3 |
| **Australia** | * | * | 98 |
| **Bahamas** | * | | |
| **Bahrain** | * | | |
| **Bangladesh** | * | | |
| **Barbados** | * | | |
| **Belize** | * | * | 64 |
| **Bermuda (UK)** | * | | |
| **Brazil** | * | * | 3 |
| **British Virgin Isl. (UK)** | * | * | 5 |
| **Brunei Darussalam** | * | | |
| **Cambodia** | * | | |
| **Cayman Isl. (UK)** | * | * | 42 |
| **Chagos Archipelago (UK)** | * | * | 49 |
| **China** | * | | |
| **Christmas Isl. (Australia)** | * | | |
| **Cocos (Keeling) Isl. (Australia)** | * | | |
| **Colombia** | * | * | 1 |
| **Comoros Isl.** | * | * | 7 |
| **Cook Islands** | * | | |
| **Costa Rica** | * | | |
| **Cuba** | * | * | 147 |
| **Djibouti** | * | | |
| **Dominica** | * | | |
| **Dominican Republic** | * | * | 36 |
| **Easter Isl. (Chile)** | * | | |
| **Ecuador** | * | | |
| **Egypt (Red Sea)** | * | * | 6 |
| **Eritrea** | * | | |
| **Fiji** | * | * | 16 |
| **French Polynesia** | * | * | 99 |

| | | | |
|---|---|---|---|
| **Grenada** | * | | |
| **Guam (USA)** | * | * | 15 |
| **Guatemala** | * | | |
| **Haiti** | * | | |
| **Hawaii** | * | * | 117 |
| **Honduras** | * | * | 3 |
| **India** | * | | |
| **Indonesia** | * | * | 132 |
| **Iran** | * | | |
| **Israel** | * | | |
| **Jamaica** | * | * | 69 |
| **Japan** | * | | |
| **Jordan** | * | | |
| **Kenya** | * | * | 27 |
| **Kiribati** | * | * | 50 |
| **Kuwait** | * | | |
| **Madagascar** | * | * | 32 |
| **Malaysia** | * | | |
| **Maldives** | * | * | 40 |
| **Marshall Isl.** | * | * | 8 |
| **Martinique (France)** | * | | |
| **Mauritius** | * | * | 7 |
| **Mayotte (France)** | * | * | 10 |
| **Mexico** | * | * | 57 |
| **Micronesia (Federated States of)** | * | * | 1 |
| **Montserrat (UK)** | * | | |
| **Mozambique** | * | * | 19 |
| **Mozambique Channel Isl. (France)** | * | | |
| **Myanmar** | * | | |
| **Nauru** | * | | |
| **Netherlands Antilles** | * | * | 31 |
| **New Caledonia (France)** | * | * | 269 |
| **Nicaragua** | * | * | 13 |
| **Niue (New Zealand)** | * | | |
| **Northern Marianas (USA)** | * | * | 24 |
| **Oman** | * | * | 7 |
| **Pakistan** | * | | |

| | | | |
|---|---|---|---|
| **Palau** | * | * | 1 |
| **Panama** | * | * | 48 |
| **Papua New Guinea** | * | * | 13 |
| **Philippines** | * | * | 1 |
| **Pitcairn (UK)** | * | | |
| **PRIA** | * | * | 128 |
| **Puerto Rico (USA)** | * | * | 23 |
| **Qatar** | * | | |
| **Reunion (France)** | * | * | 14 |
| **Saint Kitts & Nevis** | * | | |
| **Saint Lucia** | * | * | 1 |
| **Saint Vincent & the Grenadines** | * | | |
| **Samoa** | * | | |
| **Saudi Arabia** | * | | |
| **Seychelles** | * | * | 55 |
| **Singapore** | * | | |
| **Solomon Isl.** | * | * | 59 |
| **Somalia** | * | | |
| **South Africa (Indian Ocean Coast)** | * | | |
| **Sri Lanka** | * | | |
| **St Martin (France)** | * | | |
| **Sudan** | * | | |
| **Taiwan** | * | | |
| **Tanzania** | * | * | 24 |
| **Thailand** | * | | |
| **Timor Leste** | * | | |
| **Tokelau (New Zealand)** | * | | |
| **Tonga** | * | * | 6 |
| **Trinidad & Tobago** | * | | |
| **Turks & Caicos Isl. (UK)** | * | * | 20 |
| **Tuvalu** | * | | |
| **United Arab Emirates** | * | * | |
| **US Virgin Isl.** | * | * | 2 |
| **USA (Gulf of Mexico)** | * | * | |
| **Vanuatu** | * | | |
| **Venezuela** | * | * | 24 |
| **Viet Nam** | * | | |

| | | | |
|---|---|---|---|
| **Wallis & Futuna Isl. (France)** | * | * | 45 |
| **Yemen** | * | | |

**Supplementary information 1**

*Supplementary information 1.1: Sensitivity analyses to the choice of surplus production model*

We estimated sustainable reference points under three other versions of the Pella-Tomlinson model. This was done by swapping eq. 8,22 and 23 from the main manuscript with eq. S1-S4, which is known as Fletcher's re-parametrization of the Pella-Tomlinson model.

$$\mu_i = \log\left(\left({B_{0i}}^{(1-n)} + \left({B_{min}}^{(1-n)} - {B_{0i}}^{(1-n)}\right)e^{\left(\frac{yr(1-p)B_{0i}\left(\frac{1}{1+\frac{n}{2}}\right)^{\left(\frac{\frac{1}{n}+1}{2}\right)}}{B_{0i}}\right)(1-n)t}\right)^{\frac{1}{1-n}}\right) +$$

$$+\beta_5 x_{depth,i} + \beta_6 x_{crest,i} + \beta_7 x_{\frac{lagoon}{backreef},i} + \beta_8 x_{flat,i} + \beta_9 x_{pointcount,i} + \beta_{11} x_{samplingarea,i} + \beta_{12} x_{size,i} + \beta_{13} x_{grav,i} \quad \text{(S1)}$$

$$y = \frac{n^{\frac{n}{(n-1)}}}{(n-1)} \quad \text{(S2)}$$

$$MMSYi = m = rB_{0i}\left(\frac{1}{1+\frac{n}{2}}\right)^{\frac{\left(\frac{1}{n}+1\right)}{2}} \quad \text{(S3)}$$

$$B_{MMSY,i} = B_{0,i}n^{\frac{1}{1-n}} \quad \text{(S4)}$$

This model has an extra parameter, *n*, that adjusts the standing stock biomass value at which the production peaks. As we could not estimate the parameter "n" from our data, we ran four versions of it: (i) a re-parametrized version of the Gompertz-Fox model (i.e., a limiting case of the Pella-Tomlinson model as "n" approaches one; main manuscript); (ii) a Graham-Schaefer model (a version of the Pella-Tomlinson model where "n" is equal to 2; eq. S5-S7); (iii) Pella-Tomlinson with a fixed "n" equal to three; and (iii) Pella-Tomlinson with a fixed "n" equal to four (eq. S1-S4). That way we allowed the curve to peak above and below 0.5 of the estimated unfished biomass:

$$\mu_i = \log\left(\frac{B_{0,i}}{1+\left(\frac{B_{0,i}-B_{min}}{B_{min}}\right)e^{-rt_i}}\right) + \beta_5 x_{depth,i} + \beta_6 x_{crest,i} + \beta_7 x_{\frac{lagoon}{backreef},i} + \beta_8 x_{flat,i} + \beta_9 x_{pointcount,i} + \beta_{11} x_{samplingarea,i} + \beta_{12} x_{size,i} + \beta_{13} x_{grav,i} \quad \text{(S5)}$$

$$MMSYi = \frac{rB_{0,i}}{4} \quad \text{(S6)}$$

$$B_{MMSY,i} = \frac{B_{0,i}}{2} \quad \text{(S7)}$$

As expected, $B_{MMSY}$ estimates changed depending on the model used (Fig. SI1-1). However, associations with environmental factors and MMSY values remained similar to the Gompertz-Fox model.

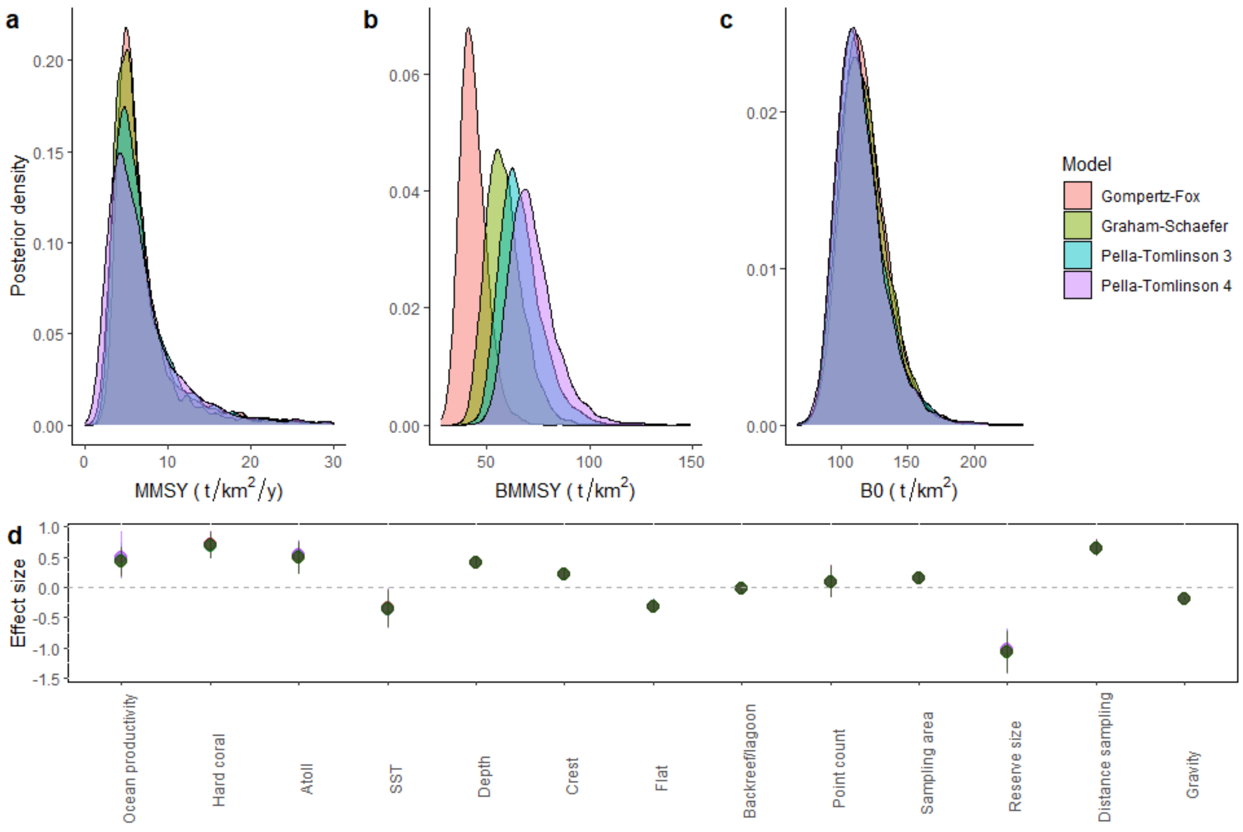

**Fig. SI1-1| Relevant reference point parameters under different surplus production models.** (a-c) MMSY, BMMSY and unfished biomass posterior distributions for average environmental conditions and non-atolls under different surplus production models. (d) Posterior effect sizes - median and 90% uncertainty intervals- of the covariates included in the reference point model, for each surplus production model.

Changing $B_{MMSY}$ values resulted in higher percentages of reefs and jurisdictions classified as below $B_{MMSY}$ and of conservation concern (Table SI1-1).

**Table SI1-1| Comparison of reference points for average environmental conditions and assessment results under different surplus production models.** Reference points are posterior medians. Percentages are using site-specific and jurisdiction median $B_{MMSY}$, MMSY and biomass estimates.

| | Gompertz-Fox | Graham-Schaefer | P-T 3 | P-T 4 |
|---|---|---|---|---|

| | | | | |
|---|---|---|---|---|
| **MMSY for average environmental conditions and non-atolls ($t/km^2/y$)** | 5.8 | 5.5 | 5.8 | 5.7 |
| **$B_{MMSY}$ for average environmental conditions and non-atolls ($t/km^2$)** | 42.6 | 57.5 | 65.0 | 70.5 |
| **% exploited reefs below $B_{MMSY}$** | 52 | 64 | 68 | 70 |
| **% jurisdictions with B below $B_{MMSY}$** | 51 | 63 | 63 | 65 |
| **% jurisdictions with $B_{weighted}$ below $B_{MMSY}$** | 37 | 47 | 55 | 59 |
| **% jurisdictions where C>MMSY** | 47 | 47 | 47 | 48 |
| **% jurisdictions of conservation concern** | 54 | 61 | 67 | 74 |

*Supplementary information 1.2: Comparison with previous reference point estimates*

Previous work estimating baselines, recovery rates and/or yields from reserve recovery trajectories using data that are also used in this study (refs (*14-15*)) have differed in best-fit parameter estimates (i.e., unfished biomass and thus $B_{MMSY}$, community growth rate, and/or MMSY), relative to the ones proposed here. For example, ref (*14* ; for the African coast), using a logistic model, provided an estimate of unfished biomass for high compliance marine reserves of ~115 t/km$^2$/y, a growth rate of ~0.23 1/y and used these (e.g*., 73, 74*) to provide seascape MMSY values of ~6.4 t/km$^2$/y. Ref (15), on the other hand, reported a global unfished baseline of ~101 t/km$^2$ and a growth rate of ~0.05 1/y. These contrast, to some degree, with our posterior medians for unfished biomass and biomass community growth rate of ~115.1 t/km$^2$ and ~0.19 1/y, respectively (Fig. S2). Here we explicitly state the differences in our approach (Table SI1-2), and we justify our model assumptions of considering open populations relative to these earlier studies.

Reserves within heavily fished seascapes are likely to export a significant portion of their production (especially if reserves are small, as in most coral reef systems; *16*). This means that (i) reserves are likely to reach asymptotic biomass values below those that would be obtained if the entire seascape was allowed to recover (and thus the reserve asymptote would be lower than the seascape unfished biomass, which is the relevant quantity for estimating MMSY and $B_{MMSY}$), and (ii) the observed per-unit-biomass recovery rates in reserves would likely be below the "true" community biomass growth rate (i.e., r), since exported growth would not be reflected in the biomass trajectory (18). As MMSY under the Gompertz-Fox surplus dynamics is equal to (r*B0)/e (where e is the Euler number), downward biasing the community growth rate and/or the unfished biomass, as would occur if reserves were assumed fully closed, would bias MMSY and $B_{MMSY}$ downward. Thus, we (i) added a human metric unstandardized to account for potential net exports from reserves, and (ii) informed the seascape-scale unfished biomass with biomass in remote locations, which represent the closest available approximation to reef systems that are unfished at the approximate scale of population closure[9], although our inclusion of environmental covariates allowed for differences in unfished biomass between reserves and remote locations due to environmental context as well. Note that we added our human impact unstandardized because adding the metric standardized (i.e., so that mathematically parameters would be estimated for average human impact) could produce a "shifting baseline" effect whereby, at average environmental conditions, locations with higher human impact would have biased reference points (e.g., $B_{MMSY}$ biased low).

Ref [1] assumed reserves acted as closed populations and did not use remote locations to inform the unfished biomass estimate from the reserve trajectory data, implicitly assuming that (i) unfished biomass was equal to asymptotic reserve biomass, and (ii) observed recovery rates to those asymptotes are equal to the community growth rate. Ref (*15*) also assumed closed populations but, in contrast to ref (*14*), did use remote locations to inform unfished baselines, thus, reserve trajectories were implicitly constrained to ultimately reach their respective remote biomass levels.

**Table SI1-2| Major differences between our approach and previous fisheries-independent studies (which data is included in this study) that estimated unfished baselines or sustainable yields for coral reef fish from reserve recovery trajectories using a logistic equation.** * indicates that the study followed the same approach as we did in our study.

| In our study | Ref *14* | Ref *15* |
|---|---|---|
| **Assumed reserves could act as open populations (including human impact)** | Assumed reserves acted as closed populations and did not include effect of human impact | |
| **Used remote reefs to inform unfished biomass** | Did not use remote reefs to inform unfished biomass | * |
| **Used space-for-time substitution for reserve recovery making sure selected sites did not impact the overall trends** | Mixed time-series and space-for-time without accounting for potential temporal autocorrelation | * |
| **Classified remote reefs systematically as >20h from human settlements, excluding locations with consistent human pressure** | NA | Classified remote reefs by expert opinion |

| | | |
|---|---|---|
| **Accounted for environmental, sampling, reserve size and the potential collinearity of these with total gravity** | Did not account for any of these effects | Accounted for some (e.g., environmental) and not others (e.g., sampling, human impact) |
| **The reference point model components included only sites that had coral cover information** | Included sites without coral cover information | * |
| **Included only species from the families in Table S1** | Included other families | |
| **Restricted the study to only tropical sites (23< abs (Lat))** | Included reef sites with larger absolute latitudes (i.e., subtropical reefs) | |
| **Global** | For the African coast | * |
| **Recovery trajectory arithmetic biomass units** | * | Recovery trajectory is log-biomass units |
| **Assumed unfished biomass varied by environmental covariates** | Assumed unfished biomass was constant (i.e., did not include environmental effects). | Assumed unfished biomass was constant (i.e., included the effect of environmental variables on observed biomass) |

*Supplementary information 1.3: Accounting for net export from reserves*

As mentioned above, reserves embedded within fished seascapes are likely exporting part of their biomass and not representing recovery at the scale of metapopulation closure. Here we explain how we attempted to parametrize exports in three different ways and what can be learned from the process.

We first parametrized exports as a proportion of the community growth rate (i.e., as r(1-p). In other words, a proportion, *p*, of the per-capita population growth was exported:

$$\frac{dB}{dt} = (\log(B0) * r * (1-p) * B * \left(1 - \frac{\log(B)}{\log(B0)}\right) \quad \text{(S8)}$$

$$\log\big(B(t)\big) = \log\left(B0 * \left(\frac{bmin}{B0}\right)^{\exp(r*t*(p\,-\,1))}\right) \quad \text{(S9)}$$

This model converged but yielded a non-identifiable export proportion (i.e., low posterior contraction: 0.22). Low posterior contraction does not necessarily indicate a problem (for example, if the prior choice is informed by theory). However, in our case we used an uninformative prior for the export proportion (uniform (0,1)), and all proportions fit our data well. Posterior samples for the export proportion were positively correlated with samples from the community growth rate such that any export value fit our data well, leading to different estimates of the community growth rate.

Next, we parametrized exports as a rate (i.e., biomass exported per biomass unit at each time step), where p has dimensions of (1/time):

$$\frac{dB}{dt} = (\log(B0) * r * B * \left(1 - \frac{\log(B)}{\log(B0)}\right) - p * B \quad \text{(S10)}$$

$$\log(B(t)) = \log\left(\exp\left(\frac{r*\log(B0) - p + \exp(-r*t)*(p - r*\log(B0) + r*\log(bmin))}{r}\right)\right) \quad (S11)$$

We tried fitting the analytical solution of this equation using a constrained export parameter such that p could not be bigger than the community growth rate r. However, this model had divergences.

We compared through leave-out-one cross validation those parameterizations and the full model employed in the main text, which allowed community growth rate and unfished biomass parameters to be estimated in the absence of human impact (i.e., zero gravity). Model selection favored the model including gravity indicating that adding such parameter increased the predictive accuracy of our model. Moreover, the gravity effect was negative, indicating that sites in higher gravity seascapes had lower biomass at a given point in time during the reserve recovery process, consistent with the idea that such locations will have more depleted biomass in the surrounding seascape, and thus will lose a larger proportion of their biomass growth to export.

Although the gravity model was favored, the next best model in terms of best predictive performance was the one that included exports as a proportion of the community growth rate (Table S2). Thus, as the export proportion had low contraction values, we then explored fixing exports to different plausible proportions (0, 5, 10, 15, 20, 25, 30%) and fitting such models to our data. This process revealed that (i) our full model including gravity was always better in terms of predictive performance, (ii) all export proportions fit our data well (expected log predictive density differences overlapped) but the model with export proportions fixed at 10 and 15% were slightly preferred out of all those fixed options (Table S2).

*Supplementary information 1.4: Species intrinsic growth rates vs community growth rates*

Estimating intrinsic growth rates for species with little or no stock assessment information, such as many coral reef fish stocks, is a challenge. However, recent efforts by 'FishLife' (*71*), have yielded rough estimates by using a combination of life-history and stock-recruitment parameters. Although some of the parameters used to estimate species level intrinsic growth rates are not well known for most coral reef fish (e.g., stock-recruitment relationship parameters) and more information is needed to understand whether reef fish follow patterns of species in better-studied systems, life-history correlates are the best available source of estimates of intrinsic growth rates for the species included in this study.

Here we provide the distribution of individual intrinsic growth rates estimated from FishLife for our reference point data weighted by abundance (to the lowest taxonomic level possible), and we overlay our estimated whole-assemblage biomass growth rate (Fig. SI1-2). Next, we estimate species intrinsic growth rates for the communities in ref (*1*) and, assuming a Graham-Schaefer model (as stated in their supporting information), overlay the implied community growth rates estimated from that study (i.e., $r=u_{mmsy}*2$). We show that it is not straightforward how individual species intrinsic growth rates translate into community growth rates (e.g., implied community growth rates can be below, above, or within the range of individual species intrinsic growth rates). There are several factors that could contribute to these patterns such as a systematic change in species composition towards, for example, slower-growing species as community biomass recovers.

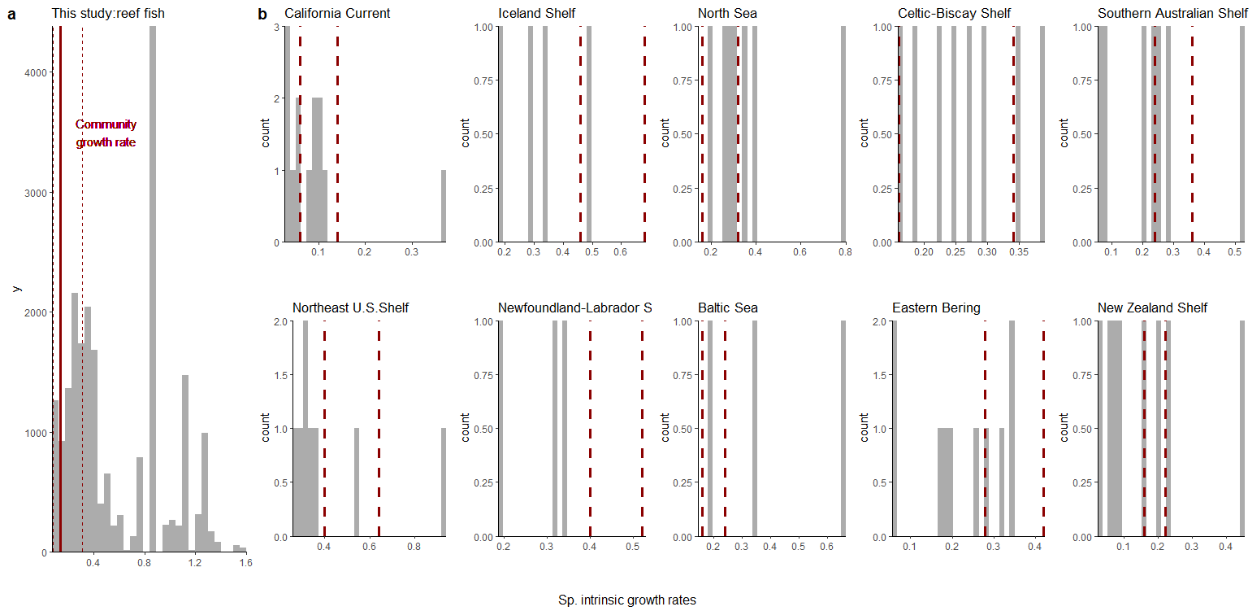

**Fig. SI1-2| Community biomass growth rate vs individual species intrinsic growth rates for reference point data (a) and the communities in ref (*1*) (b).** Histograms are the individual species intrinsic growth rates estimated from FishLife, and red dashed lines are the community growth rate intervals (90 % uncertainty intervals for (a) and reported intervals by Worm et al. 2009 assuming $r=u_{mmsy}*2$ for (b). Note that, as we had abundances for our data, in (a) histogram is weighted by abundance.

*Supplementary information 1.5: Model fits*

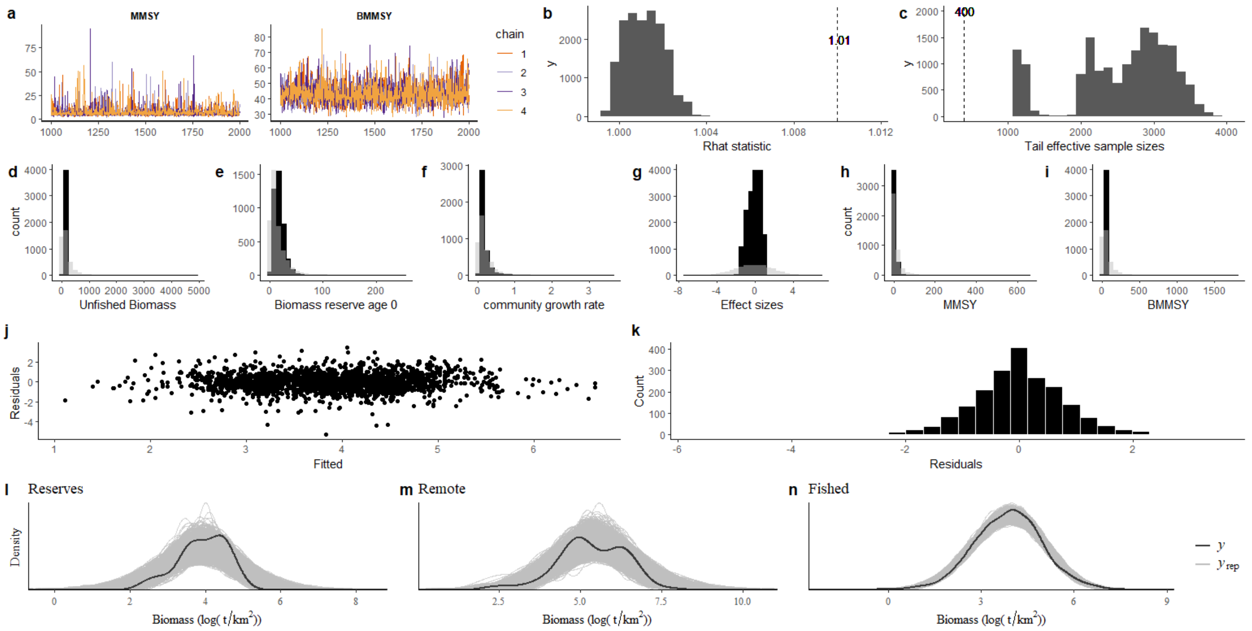

**Fig. SI1-3| Reference point model diagnostics and fit.** Example model diagnostics for benchmark model. (a) Posterior chains for MMSY and $B_{MMSY}$. (b) Potential scale reduction factor (also termed R_hat). (c) Tail effective sample sizes. (d-i) Posterior versus priors for the different estimated parameters. Light represents the prior distributions and dark represents the posterior distributions. (j) Residuals vs fitted values; (k) residual distribution; (l-n) posterior predictive checks for each model component (i.e., black line is the density of the observed data and grey lines represent the density of different simulations from the posterior predictive distribution).

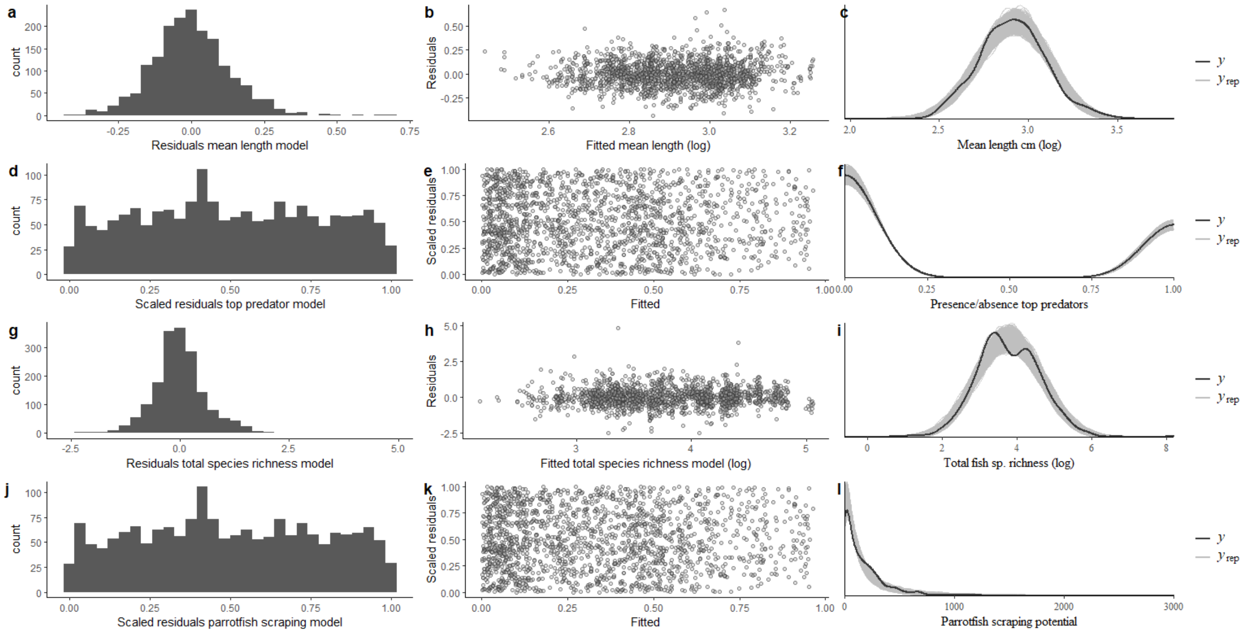

**Fig. SI1-4| Ecosystem metrics model fit.** Column 1: residual distribution; Column 2: Residuals vs fitted values; Column 3: posterior predictive checks for each model component (i.e., dark line is the density of the observed data and light lines represent the density of different simulations from the posterior predictive distribution). Note that for non-gaussian family models residuals are scaled (Dharma).

*Supplementary information 1.6: Checking data inclusion*

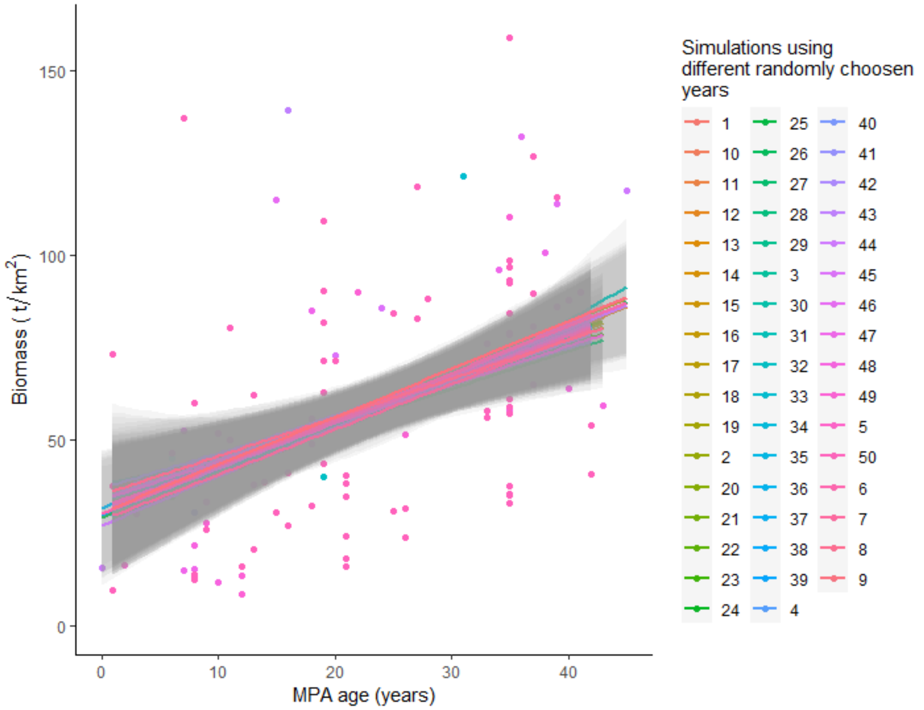

**Fig. SI1-5| Non-effect of randomly chosen reserve site years on the biomass recovery of marine reserves.** Observed biomass along a gradient of reserve ages. Model fits represent gam best-fit models with 95% confidence intervals for different randomly chosen reserve years that were duplicated (e.g., single reserve sampled multiple years).

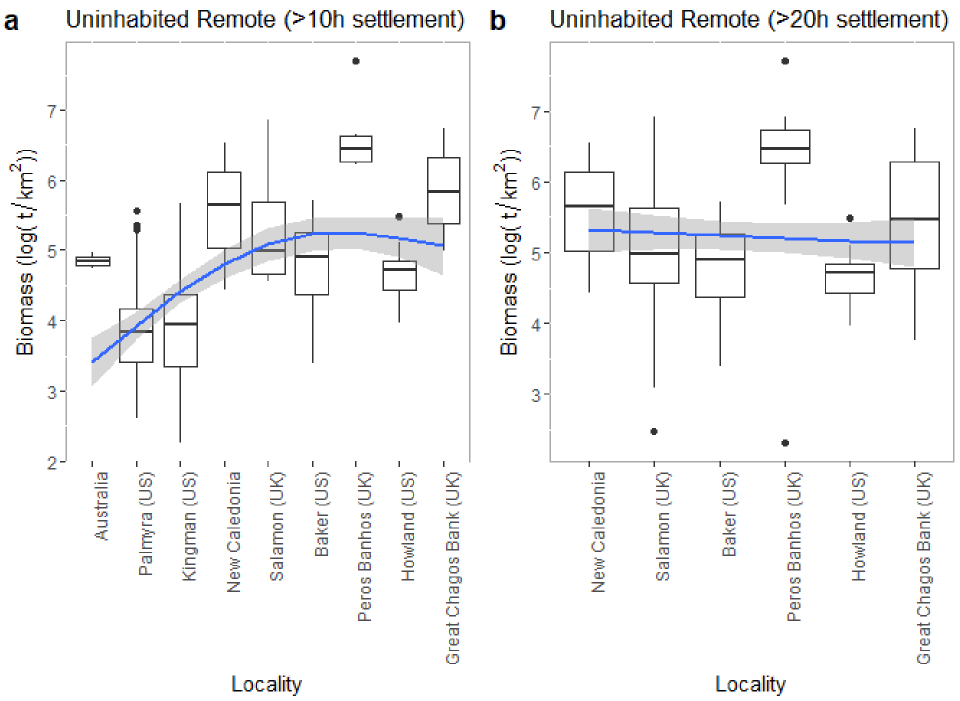

**Fig. SI1-6| Biomass in uninhabited remote reefs under different cut-off times ordered by travel time. (a) >10h away from human settlements. (b) >20 h away from human settlements.** Model fit is a generalized additive model and polygons represent the 95% confidence intervals.

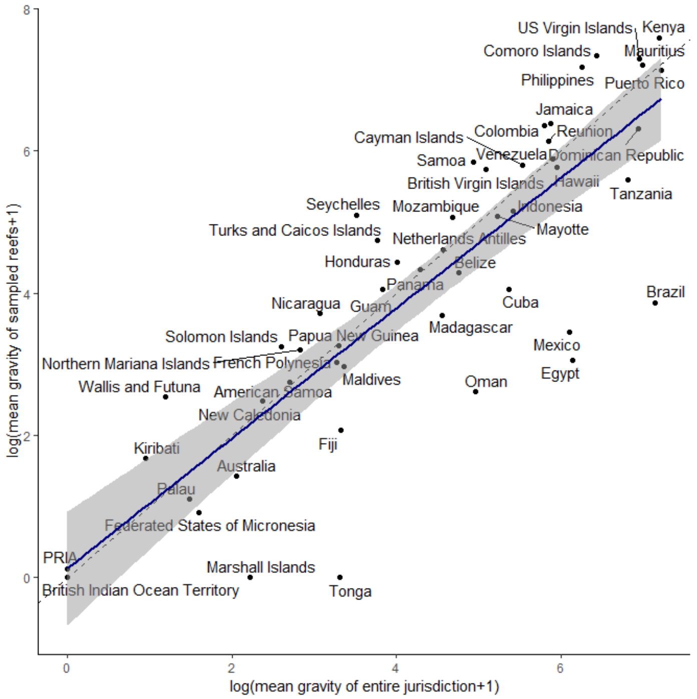

**Fig. SI1-7| Testing how representative the sampled reefs are for their jurisdiction in terms of human impact** (in a log+1 scale). Navy blue line is the linear model fit between a jurisdiction's mean total gravity and its sampled reefs mean total gravity. Polygons represent 95% confidence intervals. Dotted vertical diagonal represents the unity line.

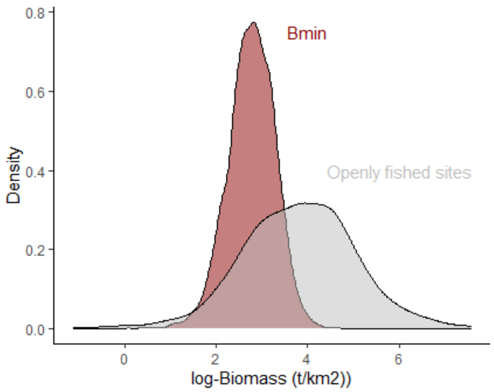

**Fig. SI1-8| Estimated Bmin vs. biomass in openly fished sites.** Densities are the posterior distribution (Bmin) and the density of observed biomass (openly fished sites).

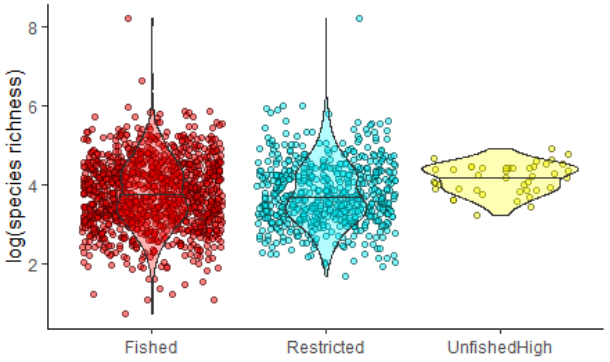

**Fig. SI1-9| Comparison of estimated fish species richness between high compliance marine reserves and exploited reefs (i.e., restricted and openly fished).** Densities show the distribution of estimated fish species richness (in log scale) separated by category. Jittered points are individual reef sites.

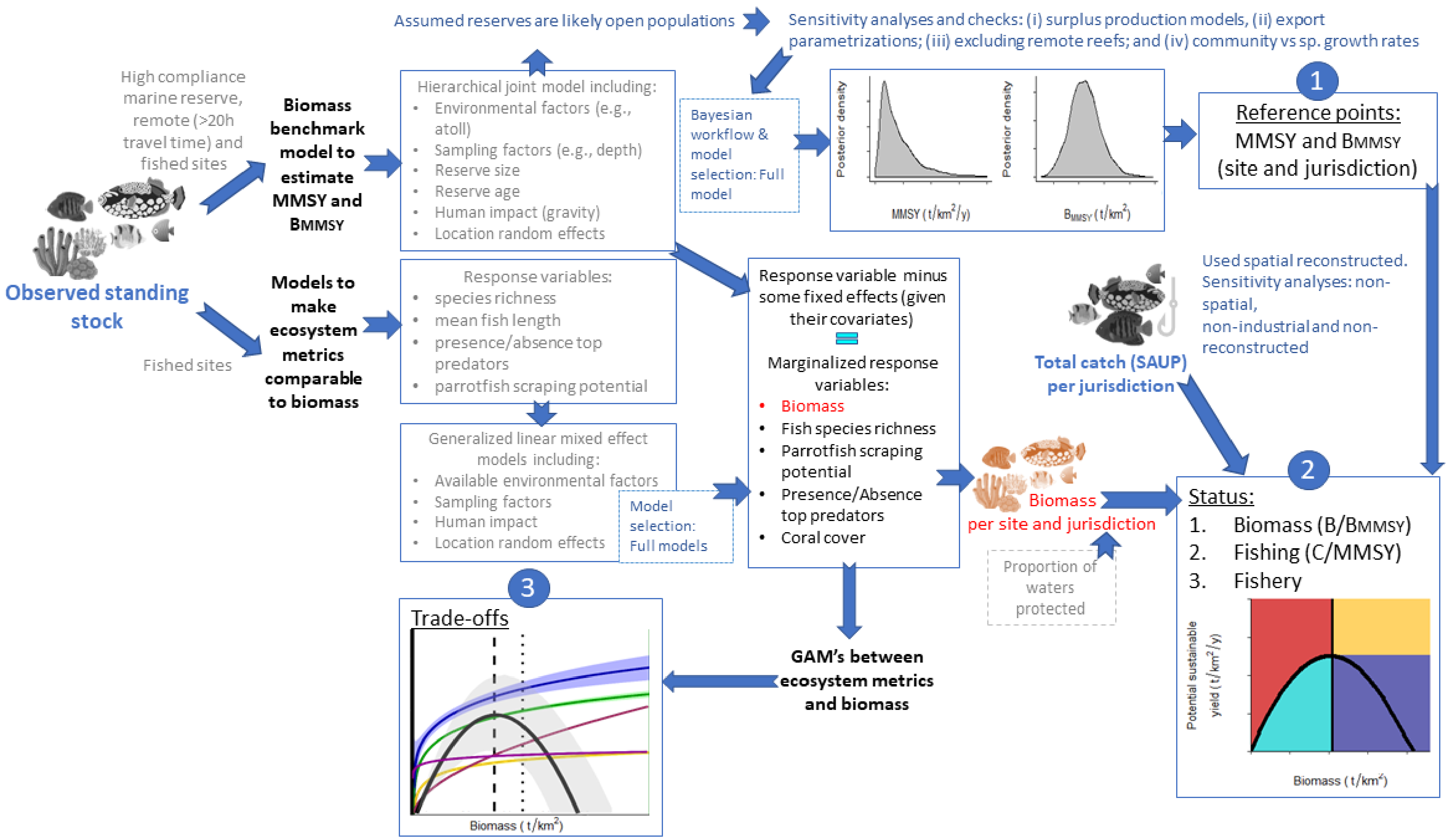

**Fig. SI1-10| Schematic of the workflow followed in this study.**

**Supplementary information 2: Principled Bayesian Workflow**

To test whether our Bayesian model accurately captured the structure of the data, we followed the Bayesian workflow in ref (*56*). First, we simulated 50 hypothetical datasets from the prior distributions and the model to check whether the simulated data are plausible and consistent with domain expertise (i.e., prior predictive checks). Secondly, we used Simulation-Based-Calibration (SBC) with the 50 simulated datasets from the prior distributions to check whether the posteriors from fitting our model to those datasets recovered the prior distributions (i.e., computational faithfulness). Thirdly, for each of the 50 posteriors, we calculated z-scores and posterior contraction values. This allowed us to test whether posteriors recover true parameters without bias and whether our model allows to identify parameters (i.e., model sensitivity). Finally, we tested whether our model is *close enough* to the true process that has generated the observed data by comparing the posterior simulations to the observed data (i.e., posterior predictive checks).

This document provides workflow results for our study and shows that the final model used (i) uses priors that are consistent with the domain expertise, (ii) produces posterior expectations that are accurate, (iii) produces posteriors that recover the true parameters (i.e., low posterior absolute z-scores) and that reduce the uncertainty of the prior (i.e., typically high contraction values), and (iv) adequately predicts the data (i.e., posterior predictive checks show posteriors overlay observed data).

The workflow was followed for two different models: the null model (without covariates or random effects), and our full model. Overall, we found that both models were good at accurately capturing the structure of the data and providing unbiased parameters (including MMSY and $B_{MMSY}$). We found that parameters estimated only from the reserve component of the model (e.g., population growth rate and biomass of reserve age zero) had, on average, good posterior contraction values, especially for the full model (also favoured in terms of predictive accuracy when fitted to our data). However, both could have low posterior contraction values for some datasets, likely indicating the relatively small number of observed data points in the reserve recovery model, in comparison with the fished component. Low posterior contraction values were more pronounced under the null model and impacted the posterior contraction of MMSY for some datasets. This highlights future research needed (i.e., more reserve data) to better inform those parameters.
Most importantly, our parameters of interest, MMSY and $B_{MMSY}$ reference points, when estimated from our data had high posterior contraction values for our final model, overall suggesting our model produces unbiased and informative MMSY reference points.
Following, we show workflow results for those models:

*Null model*

1. <u>Prior predictive checks:</u> Checking consistency of priors with domain expertise

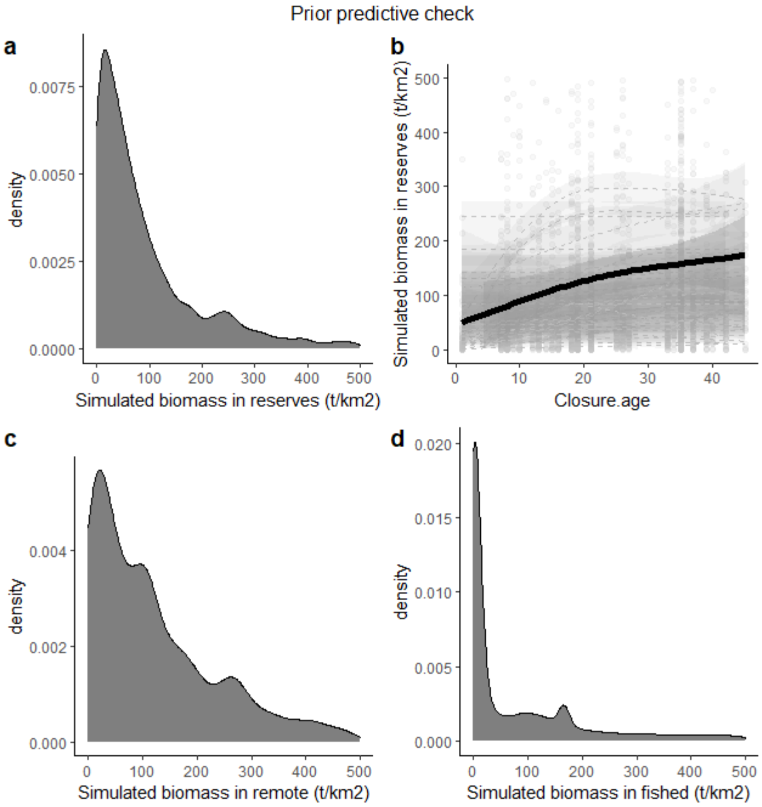

**Fig. SI2-1| Prior predictive check for our reference point null model.** (a,c,d) Combined distributions of biomass for reserves, remote and fished locations simulated from our priors. (b) Simulated biomass in reserves as a function of reserve age. Solid line is the fitted gam to the mean simulated biomass as a function of closure size. Dashed lines are the fitted gams to each individual simulation (N=50). Priors provide consistent results with our domain expertise: (i) Lognormal distributions for biomass (with most of the density at lower values); (ii) Remote reefs tend to have more biomass than fished locations; and (iii) biomass in reserves is expected to increase with reserve age.

2. Computational faithfulness: Testing for correct posterior approximations (simulation-based calibration). Checking whether posteriors from models fitted to simulated data resemble the priors used.

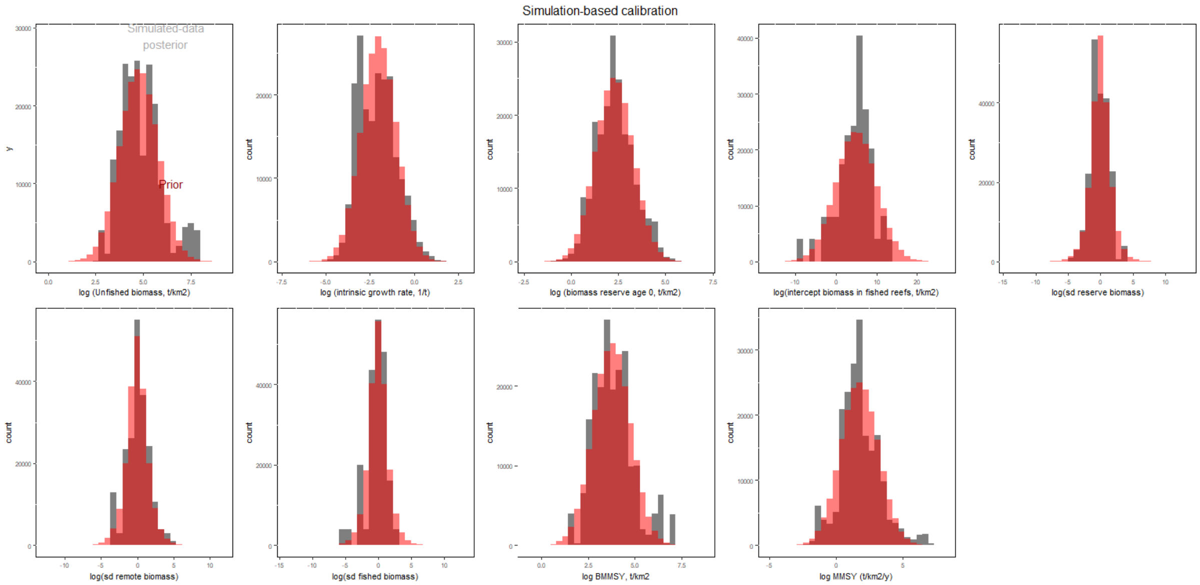

**Fig. SI2-2| Simulation-based calibration results for reference point null model**. Combined simulated data posteriors (grey) vs. distributions used to simulate the data (red) for each parameter based on simulations (N=50). Posterior expectations seem accurate for all parameters (i.e., posteriors from models fitted to simulated data overlay the prior distributions used to simulate the data).

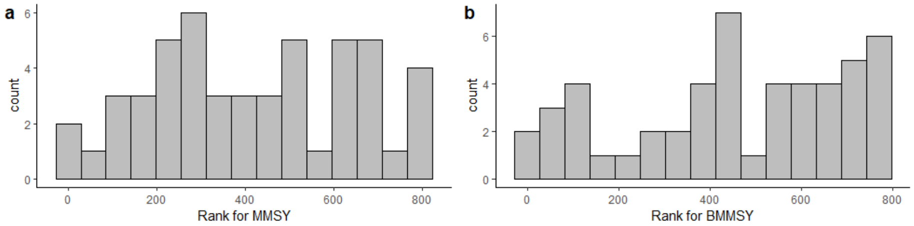

**Fig. SI2-3| Rank of MMSY and $B_{MMSY}$ based on the null model**. *"If the data averaged posterior exactly reflects the prior (identical prior and posterior; then the SBC histogram is uniformly distributed, indicating correct posterior approximation"* (ref (*56*)). Kolmogorov-Smirnov test D statistics were 0.1 for both, with bootstrap p values above 0.05 (>0.94), which means that we do not necessarily reject the null hypothesis that both distributions come from the underlying uniform distribution.

3. <u>Model sensitivity:</u> testing how well the posterior distribution answers the research question. How well does the estimated posterior mean match the true parameter used for simulating the data (z-score)? How much is uncertainty reduced from the prior to the posterior (posterior contraction)?

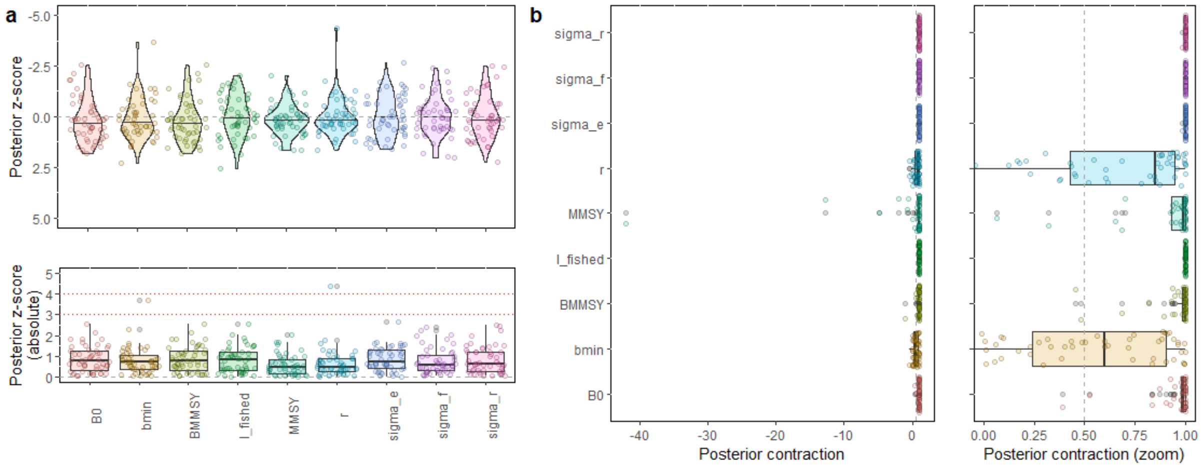

**Fig. SI2-4| Model sensitivity results for our reference point null model.** Posterior z-scores (a-raw and absolute) and posterior contraction (b-raw and zoomed) for each simulated dataset (N=50) and for each parameter. (a) The distribution of z-scores is scattered around zero, in accordance with ref (*56*): "*it is important to assess posterior z-scores for a range of simulated data sets. If no bias is present in the simulations, then the distribution of z-scores should be centered on 0, whereas shifts in the distribution of z-scores to positive or negative values indicate a bias in the posterior estimation process*." Red lines in absolute z-scores indicate the thresholds stated in ref (*56*): "*…small deviations of the estimated posterior mean are to* be expected since the posterior is fitted onto simulated data, where the simulation process will introduce some noise and thus deviations of the estimated posterior mean. Larger z-scores, e.g., larger than absolute values of 3 or 4, however, should only occur rarely due to this simulation process". (b) The majority of posterior contraction values above 0.5. However, some datasets have low posterior contraction values for the intrinsic growth rate parameter and the biomass of reserve age, creating some low posterior contraction outliers in MMSY.

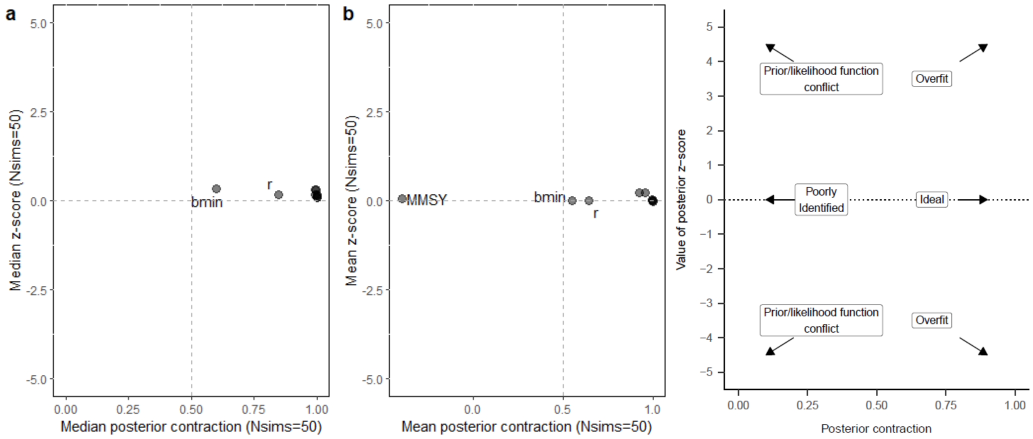

**Fig. SI2-5| Model sensitivity results for reference point null model in two dimensions.** Posterior z-scores as a function of Posterior Contraction. (a) median values based on 50 simulated datasets. (b) mean values based on 50 simulated datasets. (right) Model sensitivity classification based on ref (*56*): *"Arrows show four possible results and their interpretation. The combination of high posterior contraction with large (positive or negative) posterior z-scores reflects situations of overfitting to noise in the data. Low posterior contraction with small z-scores reflect a poorly identified model. Low contraction with large (positive or negative) z-scores indicate a substantial conflict between the prior and the likelihood. Finally, high posterior contraction and low posterior z-scores reflect an ideal situation of good model fit".* Our results suggest that our null model can provide a poorly identified MMSY parameter based on mean values (from the 50 simulated datasets), but not based on median values.

| **posterior contraction** | **parameter** |
|---|---|
| 1 | MMSY |
| 0.99 | $B_{MMSY}$ |

**Table SI2-1| Posterior contraction values for our reference points and dataset under the null model.** When fitted to our specific data, the null model produces identifiable MMSY reference points.

4. <u>Posterior predictive check:</u> Does the model adequately capture the data?
   We do this for our data. However, this is an example for one simulated dataset:

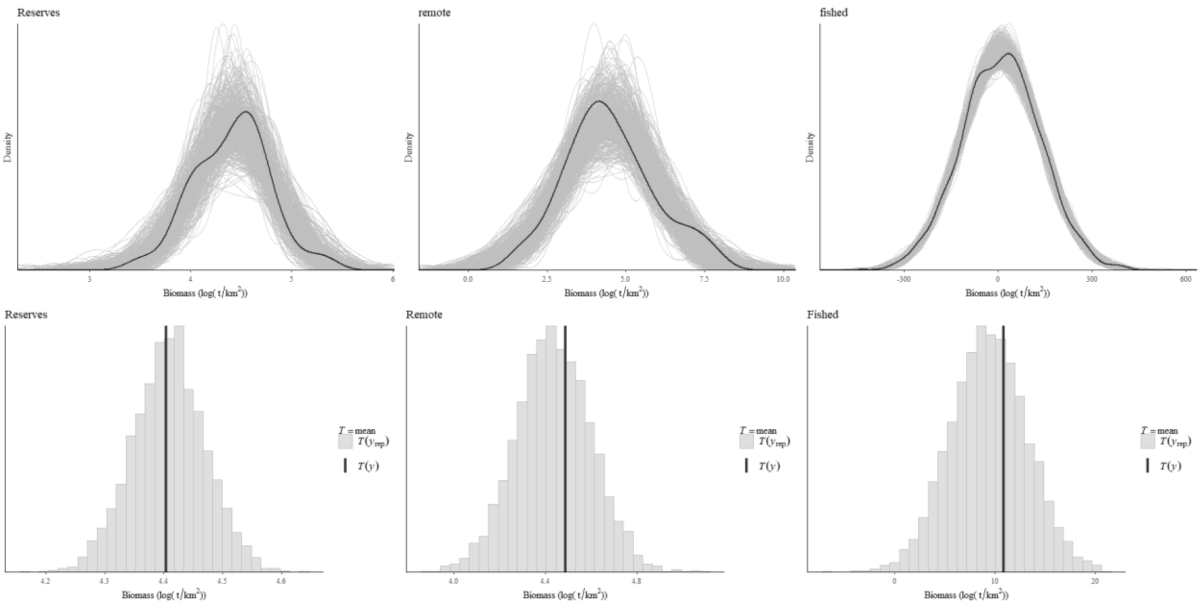

**Fig. SI2-6|Posterior predictive check for null model and simulated dataset number 50.** (top) Density of simulated data (black line) and posterior samples (grey lines). (bottom) Mean summary statistic of simulated data (black lines) and mean of the posterior samples (grey histogram). The null model captures well the simulated dataset (i.e., our model is adequate: predictions are close to the simulated data)

*Full model*

1. <u>Prior predictive checks:</u>

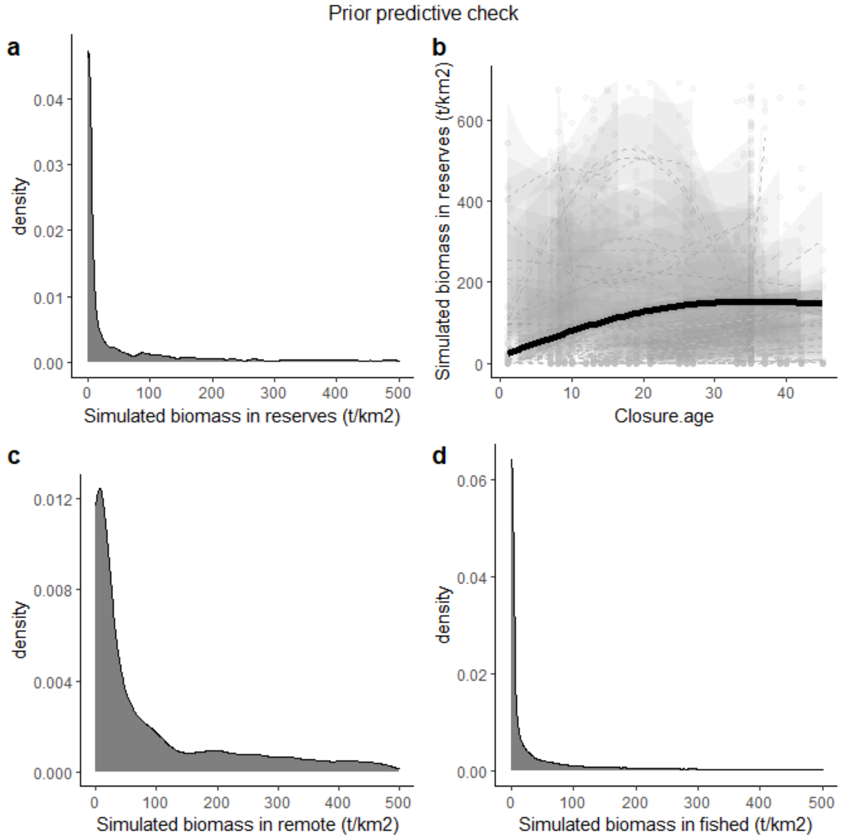

**Fig. SI2-7| Prior predictive check for our reference point full model.** (a,c,d) Combined distributions of biomass for reserves, remote and fished locations simulated from our priors. (b) Simulated biomass in reserves as a function of reserve age. Solid line is the fitted gam to the mean simulated biomass as a function of closure size (N=50). Priors fitted to our full model provide consistent results with our domain expertise: (i) Lognormal distributions for biomass (with most of the density at lower values); (ii) Remote reefs tend to have more biomass than fished locations; and (iii) biomass in reserves is expected to increase with reserve age.

2. <u>Computational faithfulness:</u>

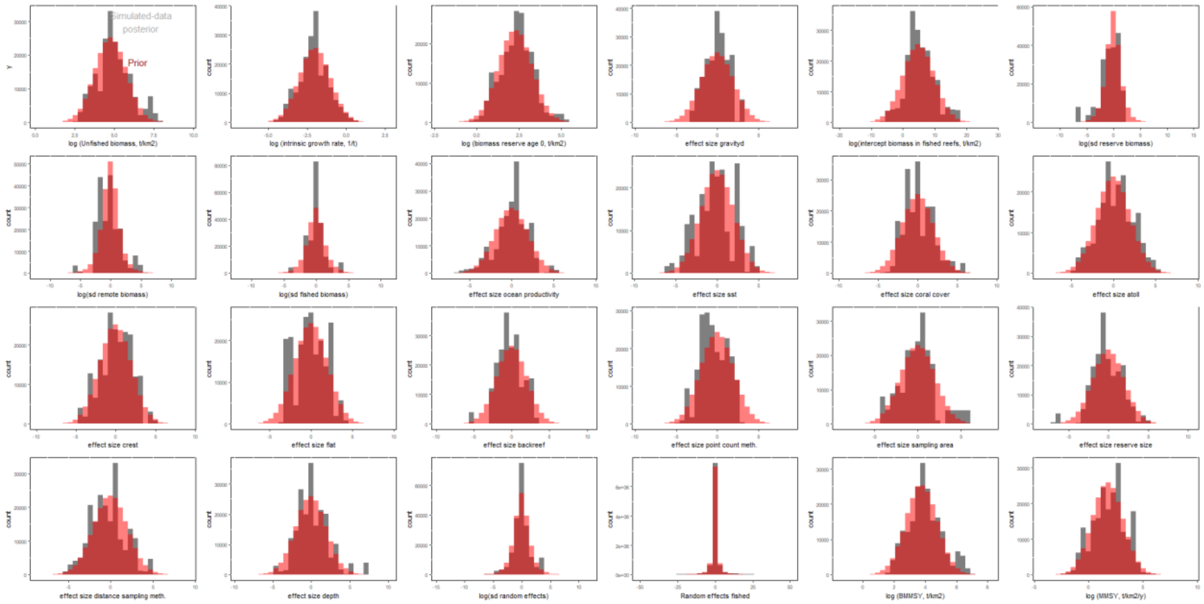

**Fig. SI2-8| Simulation-based calibration results for reference point full model.** (top) Combined simulated data posteriors (grey) vs. distributions used to simulate the data (red) for each parameter (N=50). Distributions overlay, indicating that posterior expectations seem accurate.

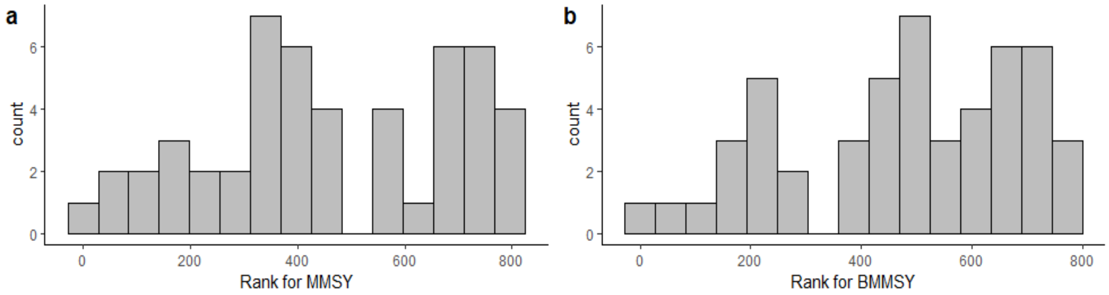

**Fig. SI2-9| Rank of MMSY and $B_{MMSY}$ based on the full model.** "*If the data averaged posterior exactly reflects the prior (identical prior and posterior; then the SBC histogram is uniformly distributed, indicating correct posterior approximation*" (ref (*56*)). Kolmogorov-Smirnov test D statistics were 0.16 and 0.14 respectively, with p values above 0.05 (> 0.51) indicating that both distributions could be samples from the uniform distribution.

3. Model sensitivity:

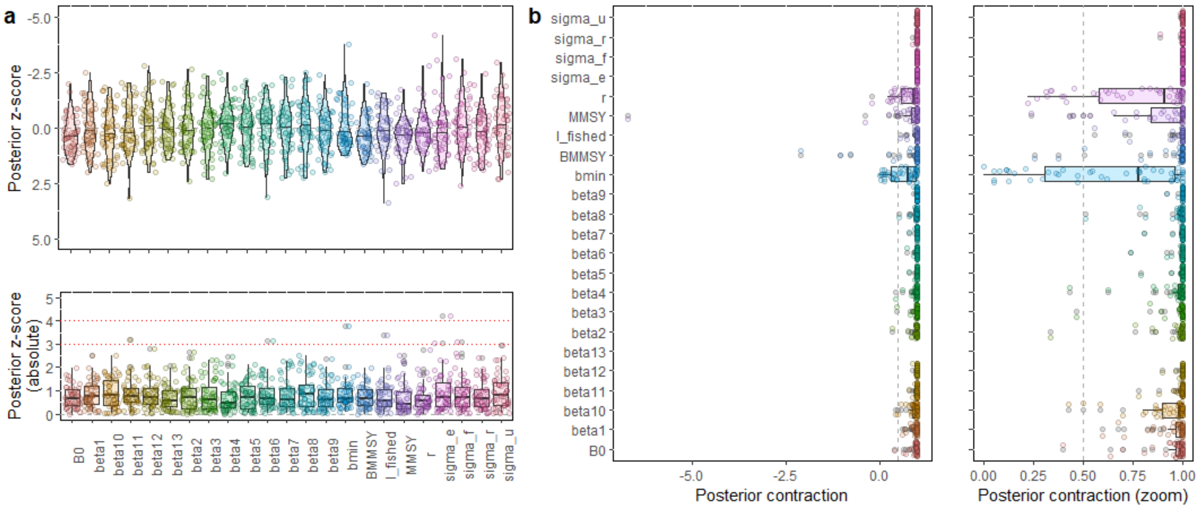

**Fig. SI2-10| Model sensitivity results for our reference point full model.** Posterior z-scores (a-raw and absolute) and posterior contraction (b-raw and zoomed) for each simulated dataset (N=50) and for each parameter. (a) The distribution of z-scores is scattered around zero, in accordance with ref (*56*): "*it is important to assess posterior z-scores for a range of simulated data sets. If no bias is present in the simulations, then the distribution of z-scores should be centered on 0, whereas shifts in the distribution of z-scores to positive or negative values indicate a bias in the posterior estimation process*." Red lines in absolute z-scores indicate the thresholds stated in ref (*56*): "*...small deviations of the estimated posterior mean are to be expected since the posterior is fitted onto simulated data, where the simulation process will introduce some noise and thus deviations of the estimated posterior mean. Larger z-scores, e.g., larger than absolute values of 3 or 4, however, should only occur rarely due to this simulation process*". (b) The majority of posterior contraction values above 0.5 (i.e., ideal zone). However, some datasets have low posterior contraction values for the intrinsic growth rate parameter and the biomass of reserve age, creating some low posterior contraction outliers in MMSY.

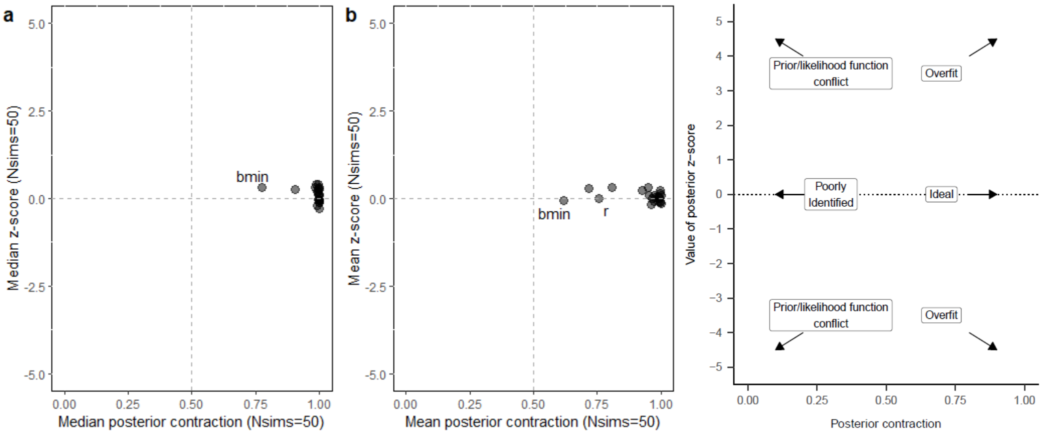

**Fig. SI2-11| Model sensitivity results for reference point full model in two dimensions.** Posterior z-scores as a function of posterior contraction. (a) median and (b) mean values based on 50 simulated datasets. (right) Model sensitivity classification based on ref (*56*): *"Arrows show four possible results and their interpretation. The combination of high posterior contraction with large (positive or negative) posterior z-scores reflects situations of overfitting to noise in the data. Low posterior contraction with small z-scores reflect a poorly identified model. Low contraction with large (positive or negative) z-scores indicate a substantial conflict between the prior and the likelihood. Finally, high posterior contraction and low posterior z-scores reflect an ideal situation of good model fit".* Our results suggest that, based on both mean and median values, our full model provides good model fit.

| posterior contraction | parameter |
|---|---|
| 0.97 | MMSY |
| 0.99 | $B_{MMSY}$ |

**Table SI2-2| Posterior contraction values for our reference point parameters and full model and fitted to our data.** Results suggest that our full model fitted to our specific data has high posterior contraction values for MMSY reference points.

4. Posterior predictive check:

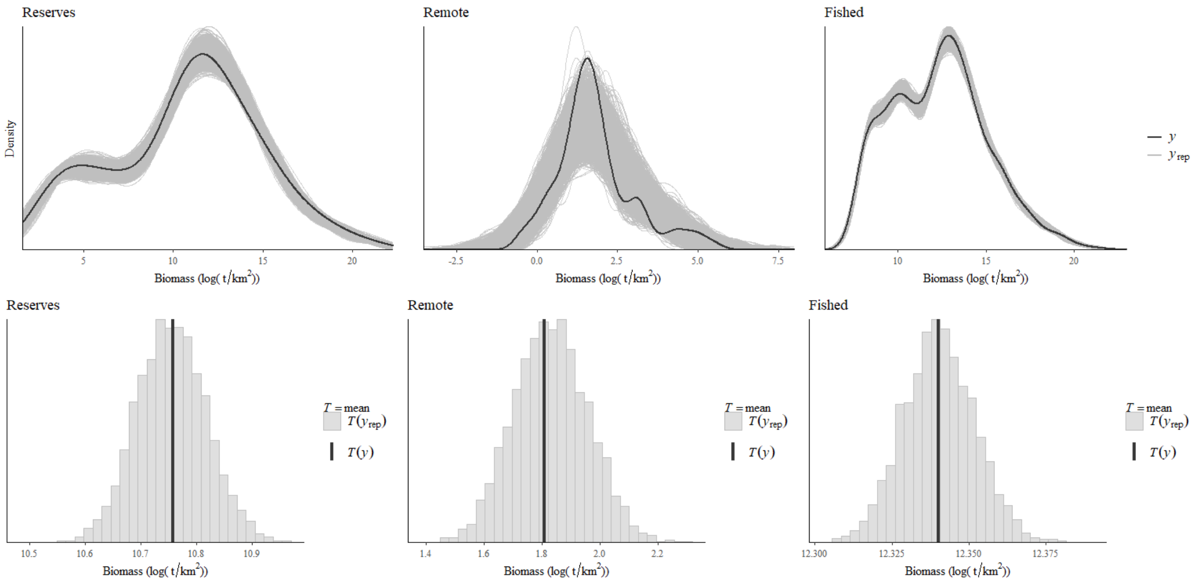

**Fig. SI2-12|Posterior predictive check for full model for simulated dataset number 50.** (top) Density of simulated data (black line) and posterior samples (grey lines). (bottom) Mean summary statistic of simulated data (black lines) and mean of the posterior samples (grey histogram). The full model fits well the data.
